# Membrane Inflammasome Activation by Choriodecidual *Ureaplasma parvum* Infection without Intra-Amniotic Infection in an NHP Model

**DOI:** 10.1101/2023.09.18.557989

**Authors:** Sudeshna Tripathy, Irina Burd, Meredith A. Kelleher

## Abstract

Intrauterine infection is a significant cause of preterm labor and neonatal morbidity and mortality. *Ureaplasma parvum* is the micro-organism most commonly isolated from cases of preterm birth and preterm premature rupture of membranes (pPROM). However, the mechanisms during the early stages of ascending reproductive tract infection that initiate maternal-fetal inflammatory pathways, preterm birth and pPROM remain poorly understood. To examine inflammation in fetal (chorioamnionic) membranes in response to *Ureaplasma parvum* infection, we utilized a novel *in vivo* non-human primate model of early choriodecidual infection. Eight chronically catheterized pregnant rhesus macaques underwent maternal-fetal catheterization surgery at 105-112 days gestation and choriodecidual inoculation with *Ureaplasma parvum* (10^5^cfu/mL of a low passaged clinical isolate, serovar 1; n=4) or saline/sterile media (Controls; n=4) starting at 115-119 days gestation, repeated every 5 days until scheduled cesarean-section at 136-140d gestation (term=167d). The average inoculation to delivery interval was 21 days and *Ureaplasma* infection of the amniotic fluid was undetectable by culture and PCR in all animals. Inflammatory mediators in amniotic fluid (AF) were assessed by Luminex, ELISA and multiplex assays. RNA was extracted from the chorion and amnionic membranes for single gene analysis (qRT-PCR) and protein expression was determined by Western blot and immunohistochemistry. Our NHP model of choriodecidual *Ureaplasma* infection, representing an early-stage ascending reproductive tract infection without microbial invasion of the amniotic cavity, resulted in increased fetal membrane protein and gene expression of MMP-9 and PTGS2, but did not result in preterm labor (no increase in uterine contractility) or increased concentrations of amniotic fluid pro-inflammatory cytokines (IL-1β, IL-6, IL-8, IL-18, TNF-α). However, membrane expression of inflammasome sensor molecules, NLRP3, NLRC4, AIM2 and NOD2, and the adaptor protein ASC (*PYCARD*) gene expression were significantly increased in the *Ureaplasma* group when compared to non-infected controls. Gene expression of *IL-1*β, *IL-18, the IL-18R1* receptor*, CASPASE-1* and pro-CASPASE-1 protein were also increased in the fetal membranes with *Ureaplasma* infection. Downstream inflammatory signaling genes MYD88 was also significantly upregulated in both the amnion and chorion, along with a significant increase in NFKB in the chorion. These results demonstrate that even at the early stages of ascending reproductive tract *Ureaplasma* infection, activation of inflammasome complexes and pathways associated with degradation of chorioamnionic membrane integrity are present. This study therefore provides experimental evidence for the importance of the early stages of ascending *Ureaplasma* infection in initiating processes of pPROM and preterm labor. These findings have implications for the identification of intrauterine inflammation before microbes are detectable in the amniotic fluid (sterile inflammation) and the timing of potential treatments for preterm labor and fetal injury caused by intrauterine infection.

## INTRODUCTION

Intrauterine infections caused by bacteria are a major cause of early spontaneous preterm birth, with 85% of births that occur earlier than 28 weeks’ gestation associated with infection and inflammation.[1] Ascending reproductive tract infection and inflammation can occur when microbes from the lower reproductive tract ascend into the choriodecidual space, where they replicate and potentially traverse intact fetal membranes to establish an intra-amniotic infection. This microbial invasion of the amniotic cavity (MIAC) is characterized, both clinically and in animal models, by a robust inflammatory response, chorioamnionitis, fetal inflammatory response syndrome (FIRS) and preterm labor.[1–3] Choriodecidual infection represents an intermediate stage in the progression of ascending reproductive tract infection prior to intra-amniotic infection and the onset of preterm labor.

*Ureaplasma* spp. are commensal bacteria of the urogenital tract and can be found in the cervix or vagina of 40-80% of sexually mature women.[4] As a species, they are the microbes most commonly associated with spontaneous preterm birth and preterm premature rupture of membranes (pPROM), being identified in 47% of placentae after preterm labor and strongly associated with histological chorioamnionitis.[5] *Ureaplasma* infections are also highly associated with neonatal morbidity, including bronchopulmonary dysplasia.[6] Furthermore, studies from a chronically catheterized maternal-fetal NHP model, have demonstrated that intra-amniotic inoculation with *U. parvum*, stimulates a robust intra-uterine pro-inflammatory response, increased uterine activity leading to premature labor, histopathological findings of chorioamnionitis and a systemic fetal inflammatory response as evidenced by fetal lung injury and pneumonitis.[3] However, this NHP model bypasses *Ureaplasma* infection of the choriodecidua prior to passage of microbes across the membranes and into the amniotic cavity. In the current study, we have therefore developed an NHP model of chronic choriodecidual infection with *Ureaplasma parvum* in order to investigate how the early stages of ascending intrauterine infection contribute to the pathogenesis of preterm labor caused by *U. parvum* infection through inflammasome activation.

Inflammasomes are multiprotein complexes that promote inflammation as part of the innate immune response through activation of the pro-inflammatory cytokines IL (interleukin)-1β and IL-18. Inflammasomes achieve this via pattern recognition receptors or “sensor” molecules, such as NLRP3 (Nucleotide-binding oligomerization domain, leucine rich repeat and pyrin domain containing-3), NLRC4 (NLR family CARD (Caspase-recruiting domain) domain-containing protein 4), NOD2 (Nucleotide-binding oligomerization domain-containing protein 2) and AIM2 (Interferon-inducible protein/Absent in melanoma 2), which respond to microbial products or tissue damage. Activation of these sensing molecules recruits adaptor proteins (ASC, apoptosis-associated speck-like protein containing a CARD, encoded by the *PYCARD* gene) and an effector protein (e.g., active forms of inflammatory caspases, CASP-1 or CASP-4), which cleave pro-IL-1β and pro-IL-18 into mature, active forms. These pro-inflammatory mediators propagate inflammation and have previously been shown to be involved in infection and inflammation associated preterm labor.[7]

The aims of this study were to determine whether localized choriodecidual *Ureaplasma parvum* infection, during an intermediate stage of ascending reproductive tract infection, is sufficient to cause activation of inflammasome complexes, and induce IL-1 *β* and IL-18 signaling in the chorioamnionic membranes, as a mechanism that may contribute to the pathogenesis of preterm labor, caused by intrauterine *Ureaplasma* infection.

## METHODS

### Animals

Studies were approved by the Institutional Animal Care and Use Committee of Oregon Health and Science University and performed in accordance with the Animal Welfare Act and Policy on Humane Care and Use of Laboratory Animals. Animals were assigned from the Oregon National Primate Research Center (ONPRC) breeding colony and ovarian hormones were measured so that animals could undergo time-mated breeding. Pregnant rhesus monkeys (*Macaca mulatta*) were then adapted to a vest and mobile catheter protection system before intra-uterine surgery was performed to implant catheters between the choriodecidua and myometrium in the lower pole of the uterus, into the amniotic cavity and maternal femoral artery and vein, as previously published [3, 8–12]. Post-operative medication included intravenous prophylactic antibiotics (Cefazolin sodium) and tocolytic medications (Terbutaline sulfate, Atosiban), as previously described.

### Experimental design

A low-passaged clinical isolate of *Ureaplasma parvum* (serovar 1) [Cassell, 1983 #1064] was provided by the Diagnostic Mycoplasma Laboratory, University of Alabama at Birmingham, USA. Animals were divided into two groups, vehicle (**Control**) and infected (***Ureaplasma, U.p.***). All animals were catheterized between 105-112 day gestational age (term=168 day). The control group (n=4) received inoculation with sterile 2SP media or saline only, and the *Ureaplasma* group (n=4) animals inoculated with *U. parvum* via indwelling choriodecidual catheters (1mL of 10^5^ CFU/mL *U. parvum* inoculum in 2SP media). Around day 140, cesarean section delivery was performed, and placenta and fetal membranes were collected and appropriately stored for downstream multiple assays. All animals were negative for *U. parvum* prior to inoculation. Infection was confirmed by quantitative culture and PCR for *U. parvum* as previously published. [3, 12, 13]

### Amniotic fluid analysis

Beginning from the time of catheterization surgery until delivery, serial amniotic fluid samples were collected via amniotic catheters. Samples were processed by centrifugation at 4400 rpm for 10 minutes at 4^°^C and the supernatant frozen at −80°C. Samples of this amniotic fluid supernatant were submitted to the ONPRC/OHSU Endocrine Technologies Core (ETC) to set up bead-based multiplex assays to analyze multiple cytokine and chemokine biomarkers using the Luminex technology (Millipore Sigma).

### Tissue collection

Immediately following cesarean section, placental tissue was collected, weights and measures recorded, with villous tissue dissected. The amnion and chorion samples were dissected from the fetal and placental side of the amniochorionic membrane, respectively. Under sterile conditions, the tissue samples were cut into small pieces, transferred to labelled cryovials, snap frozen in liquid nitrogen and stored at −80^°^C.

### RNA isolation

Total RNA was isolated from frozen amnion and chorion samples using QIAZOL^®^ Lysis reagent (Qiagen, Maryland, USA) and ZYMO Direct-zol RNA Mini Prep Plus kit (ZYMO Research, Irvine, CA, USA), according to manufacturer’s instructions. The eluted RNA was quantified using NanoDrop ND-1000 UV-VIS spectrophotometer (NanoDrop, Thermo Scientific, Wilmington, DE, USA). RNA samples with A260/A280 values of ∼1.8 −2.0 were considered as samples with negligible protein contamination.

### Real time quantitative PCR (RT-qPCR) analysis

Total RNA was reverse transcribed using Applied Biosystems^TM^ High-Capacity cDNA Reverse Transcription kit (Thermo Fisher Scientific Baltics, Lithuania), according to manufacturer’s instructions. The diluted cDNA samples equivalent to 10 ng of total RNA were subjected to validation analysis on Applied Biosystems QuantStudio 3 (Thermo Fisher Scientific) employing PowerUp^TM^ SYBR Green Master mix and 2.5 mM each of forward and reverse gene specific primers. Primers were designed from *Macaca mulatta* sequences submitted at NCBI using Primer3 and BLAST, covering the exon-exon junctions. The primer details are shown in Supplementary Table 2. Expression levels of genes were normalized to *GAPDH* expression as an internal control for each cDNA sample.

### Preparation of tissue lysates for immunoblotting

Tissue lysates were prepared using ready-to-use RIPA buffer (Sigma-Aldrich, St. Louis, MO, USA), supplemented with phosphatase inhibitor, PhosSTOP EASY pack (Roche, Mannheim, Germany) and Protease inhibitor cocktail EDTA free (Abcam). The homogenized lysates were incubated on a shaker for 30 min at 4^°^C before centrifugation at 15,000 rpm for 15 min at 4^°^C. The precipitated tissue debris pellet was discarded and the clarified lysate (supernatant) was recovered, aliquoted and stored at −80^°^C. An aliquot was subjected to colorimetric bicinchoninic acid (BCA) protein assay (Pierce, Rockford, IL, USA). Protein loading samples were prepared in Laemmli sample buffer with *β*-mercaptoethanol as reducing agent.

### Immunoblot analysis

The protein loading samples prepared in Laemmli sample buffer were resolved by Invitrogen™ Novex™ WedgeWell™ 4 to 20%, tris-glycine, 1.0 mm, mini protein gel in Novex™ tris-glycine SDS running buffer. The gel was then electroblotted onto a nitrocellulose membrane using iBlot 2 dry blotting system (Invitrogen, Thermo Fisher Scientific, Waltham, MA, USA). Non-specific sites on the membrane were blocked with 5 % milk powder in 1X TBST [10 mM Tris (pH 7.5), 150 mM NaCl and 0.05 % Tween-20] at room temperature (RT) for 1 hour and washed. The membrane was incubated overnight with primary antibody, specific for different proteins at 4°C. The membrane was incubated with loading control β-actin for 1 hour at RT and washed. The membrane was then incubated with secondary antibody (horseradish peroxide, HRP labeled anti-rabbit/ anti-mouse IgG 1:5000 diluted in 1X TBST containing 5 % milk) for 1 hour at RT. At the end of the incubation, membrane was washed and developed using a SuperSignal^TM^ west pico plus chemiluminescent substrate (Thermo Scientific, IL, USA). The blot was developed in iBright 1500 imaging system (Invitrogen, Thermo Fisher Scientific, Waltham, MA, USA) and quantitated by densitometry using ImageJ (NIH). Primary antibodies used in the present study are detailed in Table 1.

**Table 1.** List of Antibodies Used.

| Antibody # | Protein symbol | Protein name | Company | Catalog # |
| --- | --- | --- | --- | --- |
| 1 | NLRP3 | NLRP3 (D4D8T) | Cell Signaling Technology | 15101 |
| 2 | NLRC4 | NLRC4 (D5Y8E) | Cell Signaling Technology | 12421 |
| 3 | AIM2 | AIM2 (D5X7K) | Cell Signaling Technology | 12948 |
| 4 | NOD2 | NOD2 (2D9) | Gene Tex | GTX30615 |
| 5 | ASC/TMS1 | ASC/TMS1 (E1E3I) | Cell Signaling Technology | 13833 |
| 6 | IL-18 | Human/Rhesus Macaque IL-18/IL-1F4 Antibody | R&D Systems | AF2548 |
| 7 | CASPASE1 | Recombinant Anti-pro Caspase-1 + p10 + p12 antibody [EPR16883] | Abcam | ab179515 |
| 8 | IL-1β | IL-1 beta/IL-1F2 | Novus Biologicals | NB600-633 |
| 9 | PTGS2/COX2 | Recombinant Anti-COX2 / Cyclooxygenase 2 | Abcam | ab179800 |
| 10 | MM9 | Anti-MMP9 antibody | Abcam | ab58803 |
| 11 | ACTB | Beta-Actin Monoclonal Antibody (15G5A11/E2) | Invitrogen | MA1-140 |

### Immunohistochemistry

After cesarean section, fetal membrane roll was fixed in 10% neutral buffered formalin and paraffin embedded. Histological sections were cut at 5 µm thickness and mounted on slides. The slides were stained with hematoxylin and eosin (H&E), observed and photographed by Carl Zeiss Axio Scope. The H&E-stained sections were examined by a placental pathologist (TKM) blinded to treatment group for characteristics of chorioamnionitis and immune cell infiltrates. The unstained histology slides were deparaffinized and hydrated with a series of histo-clear (National diagnostics) and ethanol washes. The antigen retrieval was performed either in Sodium citrate buffer or Tris-EDTA buffer (10 mM Tris-base, 1 mM EDTA solution, 0.05 % Tween 20, pH 9.0), according to primary antibody manufacturer’s instructions under high pressure. After blocking the non-specific binding with VECTASTAIN Elite ABC HRP Kit (Peroxidase, Anti-Mouse/Anti-Rabbit IgG) Kit (Vector Laboratories, CA, USA), the sections were incubated overnight with non-fluorophores primary antibody at 4^°^C. Secondary antibody incubation was followed by the ABC reagent from VECTASTAIN Elite ABC HRP Kit. The sections were developed using ImmPACT DAB Peroxidase (HRP) Substrate Kit (Vector Laboratories, CA, USA), mentioned earlier. Slides were mounted and imaged using the Carl Zeiss Axio Scope with AxioCam ERc 5s. The details of primary antibodies used in the present study are put in Table 1.

### Statistical Analyses

Data is expressed as mean±SEM unless indicated. Normal distribution was determined using Shapiro-Wilk test. Comparisons between two groups were carried out using a Students’ T-Test for normally distributed data or Mann-Whitney U-test for non-parametric data. A p-value of <0.05 was considered to be significant. All data were analyzed using statistical analysis software (GraphPad Prism, La Jolla, CA).

## RESULTS

### Ureaplasma parvum Infection did not spread from the Chorioamnionic Membranes following Choriodecidual Inoculation

Animals in both control and *U.p.* groups had negative amniotic fluid cultures indicating that microbial invasion of the amniotic cavity did not occur for the length of the experiments (Supplementary Table 1). For all animals in *U.p.* group, swabs collected from the choriodecidual inoculation site at the time of C-section were positive for live *Ureaplasma parvum* by culture and PCR. Two of four animals (U.p. 6 and U.p. 8) in the *Ureaplasma* group were also positive by culture and PCR in swabs of the fetal membranes, distal from the inoculation site and in placental tissue samples. A third animal in the *U.p.* group (U.p. 5) was positive by PCR (but not culture) for *Ureaplasma* present in the membranes, distal from the inoculation site. The fourth animal (U.p. 7) in the *U.p.* was negative by culture and PCR for samples other than directly at the inoculation site.

### Choriodecidual Ureaplasma Parvum Infection does not induce Preterm Labor or Intra-Amniotic Inflammation

There were no significant differences in pregnancy outcomes or gestational age at the time of surgical procedures of C-section delivery between animals in the control or *Ureaplasma* groups. Animals in the *Ureaplasma* group were exposed to choriodecidual infection for an average period of 20 days (Table 2) from the time of the first inoculation until C-section delivery at 136-140 days gestation. Uterine activity was similar between animals in the control and *Ureaplasma* animal groups, both prior and following choriodecidual inoculation (Table 2). Animals in both groups did not go into preterm labor (defined by increased uterine activity) prior to the scheduled C-section. Fetal sex was evenly distributed in each group (50% male/female) and no gross fetal abnormalities were identified during necropsy or tissue collection.

**Table 2.** Pregnancy Outcomes, Animal Characteristics, Uterine Contractility and Amniotic Fluid Inflammatory Mediators in Control and Choriodecidual *Ureaplasma* Infected Animals.

|  | Vehicle Control (n=4) | *Ureaplasma* (n=4) | P-Value |
| --- | --- | --- | --- |
| **MATERNAL & FETAL OUTCOMES (days)** |  |  |  |
| Gestational Age at Catheter Surgery | 107 (1.8) | 109 (0.5) | 0.23 |
| Gestational Age at 1st inoculation | 116 (0.9) | 119 (0.3) | 0.09 |
| Inoculation to Delivery Interval | 23 (1.3) | 20 (0.5) | 0.07 |
| Gestational Age at C-Section | 139 (1) | 138 (0.6) | 0.49 |
| Fetal Sex: Male (Female) | 2 (2) | 2 (2) | 0.50 |
| **UTERINE CONTRACTILITY (mmHg.s/hr)** |  |  |  |
| Mean Hourly Integral | 1083 (81.4) | 1052 (214.5) | 0.90 |
| Peak Hourly Integral | 3267 (545.9) | 3618 (1172) | 0.80 |
| **AMNIOTIC FLUID INFLAMMATORY MEDIATORS** |  |  |  |
| IL-1β (pg/mL) | 0.7 (0.7) | 0 (0) | 0.50 |
| TNF-α (pg/mL) | 1.6 (1.2) | 3 (2.5) | 0.41 |
| IL-6 (ng/mL) | 2.7 (1.3) | 4.2 (2.1) | 0.27 |
| IL-8 (ng/mL) | 2.7 (1.4) | 1.7 (4.4) | 0.27 |
| IL-18 (pg/mL) | 4 (4) | 0 (0) | 0.50 |
| PGE2 (pg/mL) | 75.4 (12.9) | 92.1 (23.6) | 0.28 |
| PGF2α (pg/mL) | 132.1 (28.6) | 87.8 (15.2) | 0.11 |
| MMP-1 (ng/mL) | 6.7 (1.5) | 4.5 (0.9) | **0.04\*** |
| MMP-3 (ng/mL) | 1.4 (0.3) | 1.3 (0.2) | 0.42 |
| MMP-9 (pg/mL) | 184.8 (0) | 184.8 (0) | 0.50 |
| MMP-13 (pg/mL) | 153.8 (0.8) | 162.2 (2.6) | **0.04\*** |
| Leukocytes (cells/μL) | 1.3 (0.2) | 10 (2.6) | **0.02\*** |

We identified a statistically significant increase in the peak post-inoculation number of leukocytes present in the amniotic fluid of *U. parvum* infected animals compared to control (10.6 VS 1.3 cells/uL; P=0.038). However, this is a relatively low absolute number of amniotic fluid leukocytes compared to those present with direct intra-amniotic infection with the same strain of *U. parvum* in rhesus monkeys.[11] Inflammatory mediators in the amniotic fluid were compared across serial samples taken prior to and following choriodecidual inoculation. Post-inoculation peak values for mediators commonly associated with inflammation-mediated preterm labor (IL-1*β*, TNF-*α*, IL-6, IL-8, and IL-18; Table 2) were not significantly different between control and *Ureaplasma* infected animals. There was no significant elevation between baseline (pre-inoculation) and post-inoculation concentrations when compared within and across groups (2-way ANOVA; *P*>0.05) for all mediators measured, indicating minimal inflammation within the amniotic cavity.

### Increased Gene and Protein Expression of Inflammasome Sensor Molecules in the Chorioamnionic Membranes Exposed to Choriodecidual Ureaplasma Parvum Infection

The expression of major inflammasome sensor molecules (NLRP3, NLRC4, AIM2, NOD2) were determined in the amnion and chorion. In the amnion, there was a significant upregulation in the mRNA expression of *NLRC4* in the *U. parvum* animals, compared to control (P<0.05), but no significant changes in *NLRP3*, *AIM2* or *NOD2* gene expression (Fig. 1A). Western blot analysis demonstrated increased protein expression of AIM2 in the amnion (Fig. 1B&C). When examined in the chorion, NOD2 gene expression was upregulated in the *U.p.* group (Fig. 2A, P<0.05). *NLRP3* gene expression was increased (P=0.07, Fig. 2A), accompanied by significantly increased NLRP3 protein expression (P<0.05, Fig. 2B&C) in the chorion of animals with choriodecidual *U. parvum* infected animals compared to controls.

**Figure 1.**
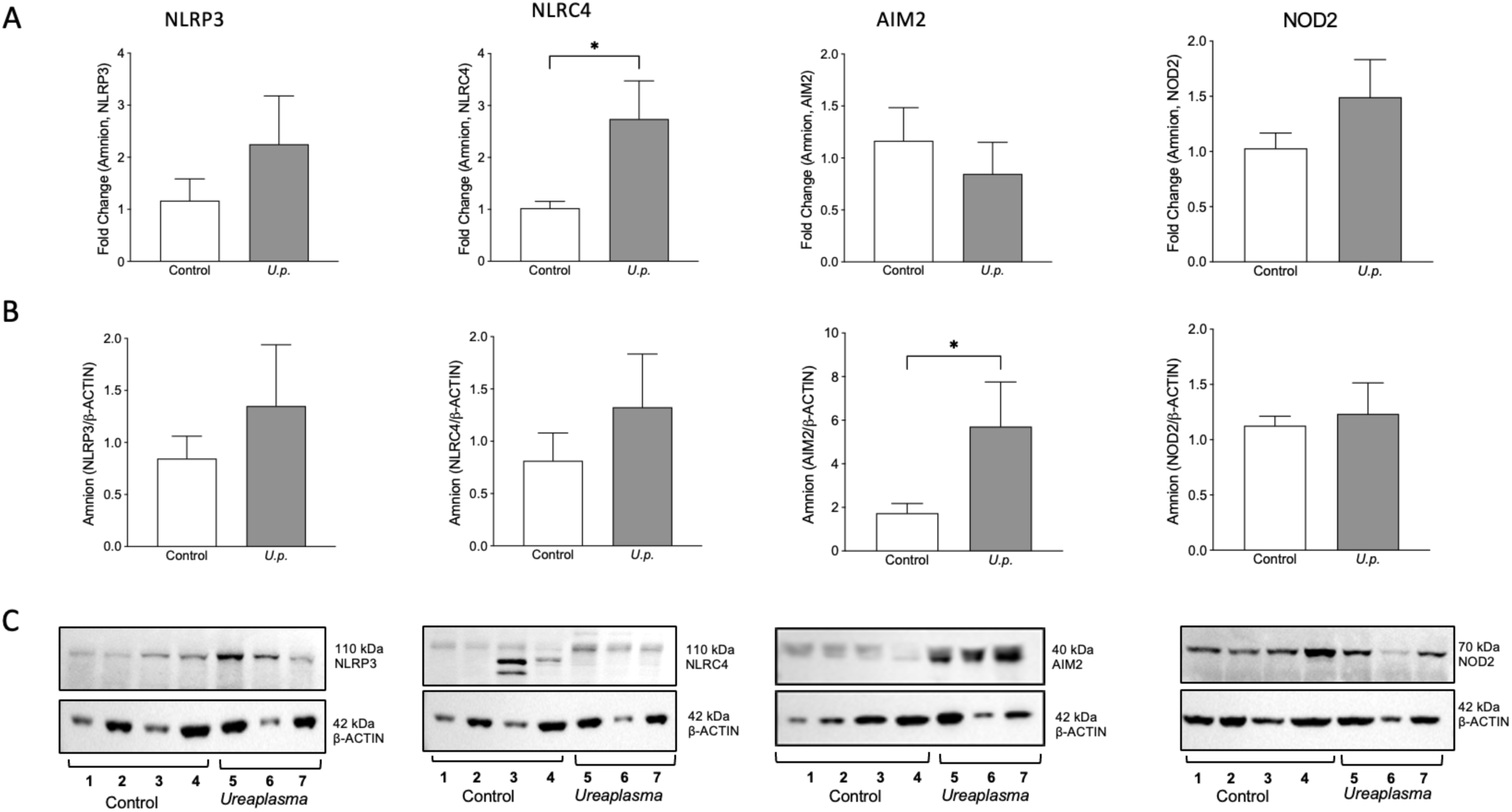
Choriodecidual *Ureaplasma parvum* infection induces an increase gene and protein expression of inflammasome sensor molecules in amnion. (A) q(RT)PCR expression of *NLRP3*, *NLRC4*, *AIM2* and *NOD2* in the amnion collected following preterm Cesarean section delivery. The results are shown as fold changes in mRNA expression compared with control group. *GAPDH* is used as the housekeeping gene. Individual bars represent mean±SEM fold change per group. The asterisks indicate significance difference from the corresponding control. (B) Densitometric quantification and (C) representative western blot of NLRP3, NLRC4, AIM2 and NOD2 protein expression in the amnion from control and choriodecidual *Ureaplasma parvum (U.p.)* infection group. *β*-actin serves as a loading control. The results are shown as relative protein levels with respect to the loading control. Each bar represents the mean±SEM. The asterisks indicate significance difference from the corresponding control (Student’s t-test/ Mann-Whitney U-test, *P<0.05, n=3-4 animals/group).

**Figure 2.**
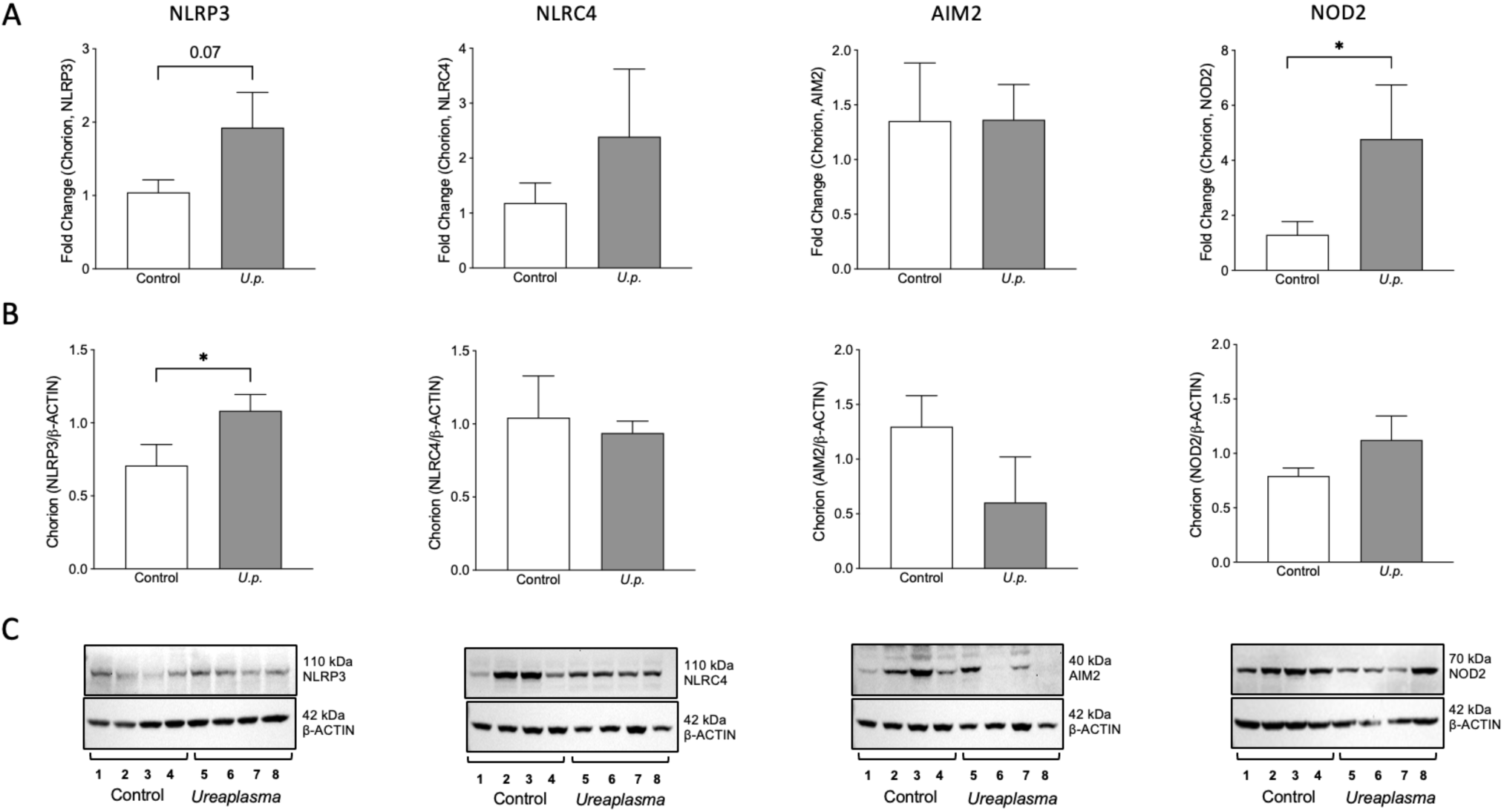
Choriodecidual *Ureaplasma parvum* infection induces changes in gene and protein expression of inflammasome sensor molecules in chorion. (A) q(RT)PCR expression of *NLRP3*, *NLRC4*, *AIM2* and *NOD2* in the chorion collected following preterm cesarean section delivery. The results are shown as fold changes in mRNA expression compared with control group. *GAPDH* is used as the housekeeping gene. Individual bars represent mean±SEM fold change. The asterisks indicate significance difference from the corresponding control. (B) Densitometric quantification and (C) representative western blot of NLRP3, NLRC4, AIM2 and NOD2 in the chorion from control and choriodecidual *Ureaplasma parvum (U.p.)* infection group, as indicated. *β*-actin serves as a loading control. The results are shown as relative protein levels with respect to the loading control. Each bar represents the mean± SEM. The asterisks indicate significance difference from the corresponding control (Student’s t-test/ Mann-Whitney U-test, *P<0.05, n=4 animals/group).

### Increased PYCARD and CASPASE-1 mRNA Expression in Chorioamnionic Membranes

Expression of *PYCARD* mRNA (Fig. 3A&B), the gene that encodes the ASC adaptor protein of the inflammasome complex, was significantly increased in the chorion of animals exposed to choriodecidual *Ureaplasma* infection (P<0.05, Fig. 3B). *PYCARD* mRNA in the amnion and ASC protein expression in both the amnion and chorion remained similar between the control and choriodecidual infection groups (Fig. 3A&B).

**Figure 3.**
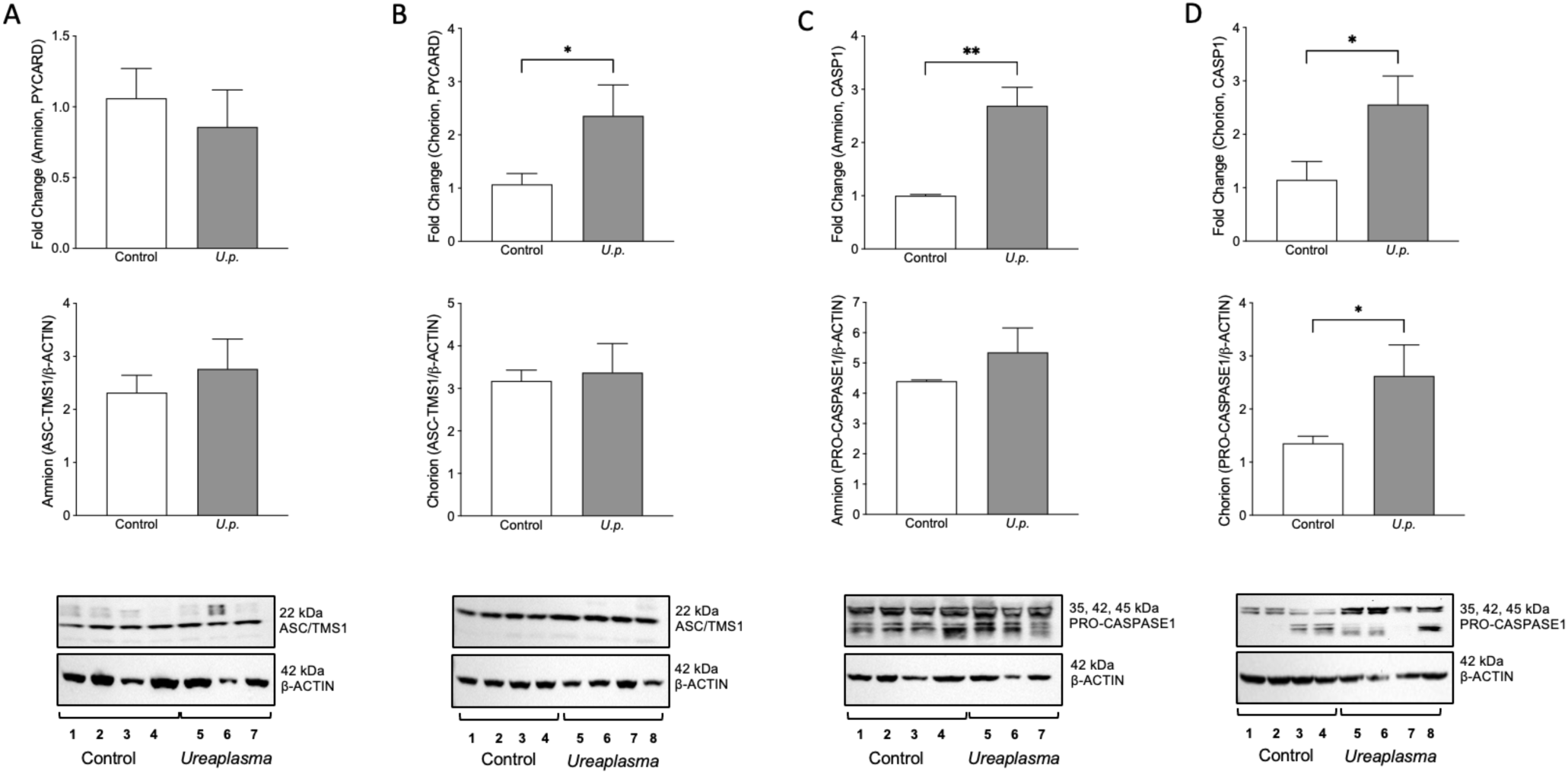
Choriodecidual *Ureaplasma parvum* infection induces *PYCARD (ASC)* and *CASP1* gene expression in chorioamnionic membranes. For the adaptor component of the inflammasome complex, q(RT)PCR expression of the *PYCARD* gene and corresponding ASC protein expression and representative Western blots are shown in the (A) amnion and (B) chorion. For the effector component of the inflammasome complex, q(RT)PCR expression of *CASP1* gene and corresponding pro-CASPASE-1 protein expression and representative Western blots are shown in the (C) amnion and (D) chorion. The results are shown as fold changes in mRNA expression compared with control group and *GAPDH* as the housekeeping gene. For protein, results are shown as relative protein expression with respect to the *β*-actin loading control. Individual bars represent mean±SEM fold change. The asterisks indicate significance difference from the corresponding control (Student’s t-test/ Mann-Whitney U-test, *P<0.05, **P<0.01, n=3-4 animals/group).

The expression of the inflammasome effector protein, *CASP-1* mRNA was significantly upregulated in both the amnion and chorion in the *U.p.* group animals when compared to controls (P<0.05, Fig. 3C&D). When CASPASE-1 protein was measured by Western Blot, neither the control or Ureaplasma groups expressed bands at the expected size for mature CASP-1 (Fig. 3C&D). However, a significant upregulation in the immature, uncleaved pro-CASPASE-1 protein was identified in the chorion membranes of the infection group (Fig. 3D).

### Pro-inflammatory IL-18, IL-1 α and IL-1 β mRNA Expression Upregulated in Chorioamnionic Membranes following Choriodecidual Ureaplasma parvum Infection

Pro-Inflammatory genes IL-8 and TNF-*α* were upregulated with choriodecidual *Ureaplasma* infection in the amnion (Fig.4A) and chorion (Fig. 4B), respectively. Gene expression of members of the pro-inflammatory IL-1 family cytokines (including IL-18) that can be activated by inflammasome action and their receptors were measured in the amnion (Fig. 4A) and chorion (Fig. 4B). mRNA for the cytokines *IL-1α* and *IL-1β* were significantly upregulated in the amnion (P<0.05) in *U.p.* animals compared to controls. The IL-18 gene expression was significantly upregulated in both amnion and chorion. The gene for the IL-18 receptor 1 (*IL-18R1)* was also significantly increased in the amnion. Gene expression of *IL-18* and *IL-18BP* were significantly increased in the chorion (P<0.05, Fig. 4B). *IL-1β* and IL-18 protein expression were also measured by Western blot in order to determine expression of the pro- and the mature forms that can be activated by CASPASE-1. Relative concentrations of Pro-IL-18 and mature IL-18 did not differ between control and *U. parvum* groups in the amnion or chorion (supplementary Fig. 1). The ratio of Mature/Pro IL-18 was also similar in the amnion or chorion between the groups. Only pro-*IL-1β* protein at 31kDa was detected in the amnion and chorion membranes and no changes in expression were identified between the two groups (supplementary Fig. 2).

**Figure 4.**
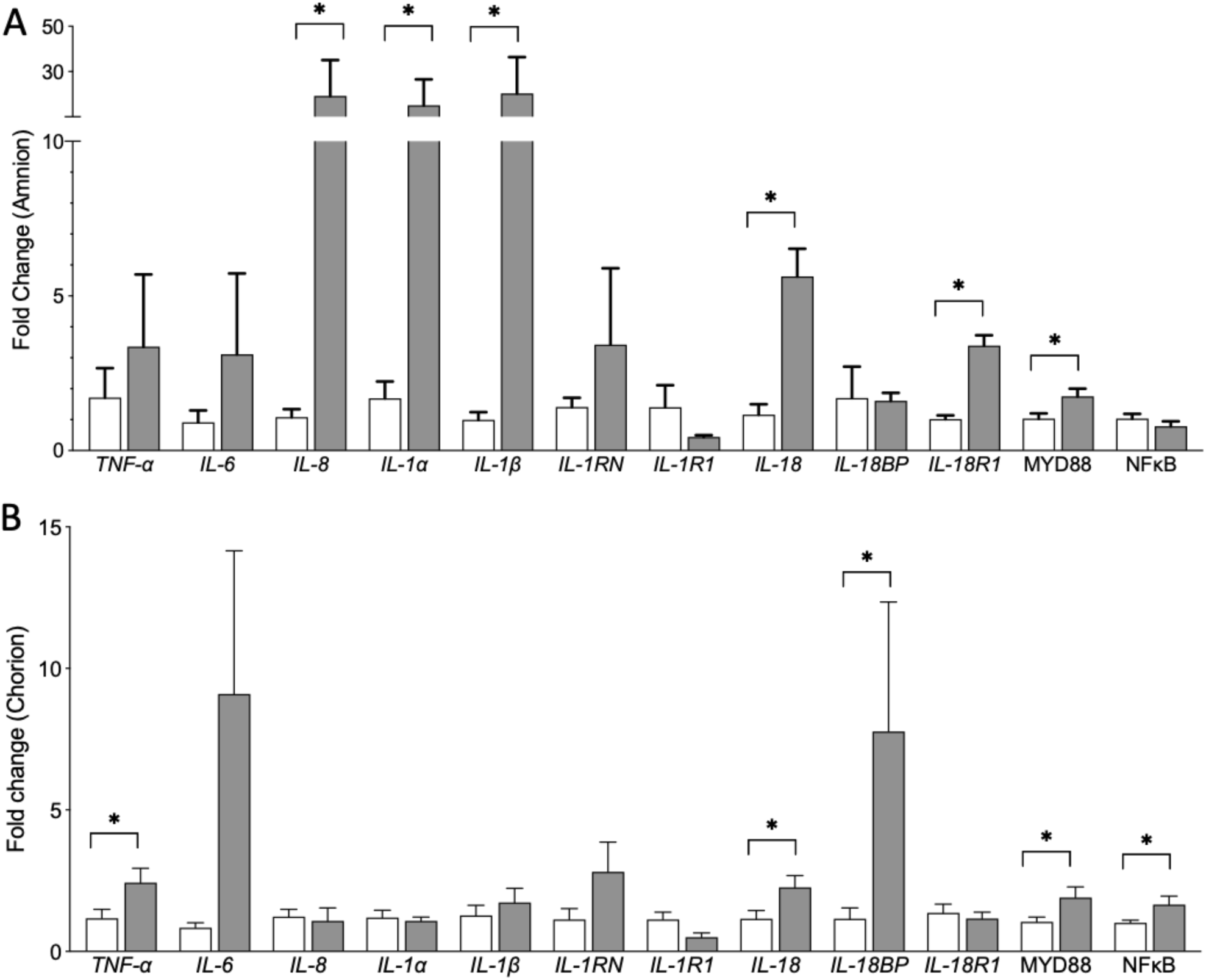
Choriodecidual *Ureaplasma parvum* infection induces an increase in gene expression of *IL-1* family members and *IL-18* in chorioamnionic membranes. q(RT)PCR expression of *TNF-⍺*, *IL-6*, *IL-8*, *IL-1⍺*, *IL-1β*, *IL-1RN*, *IL-1R1*, *IL-18*, *IL-18BP* and *IL-18R1* in the (A) amnion and (B) chorion collected during cesarean section delivery. Downstream gene expression of *MYD88* and *NFκB* are also shown. The results are expressed as fold changes in mRNA expression compared with control group and the housekeeping gene *GAPDH*. Individual bars represent mean±SEM fold change. Open bars are control and grey bars are the *Ureaplasma* group. The asterisks indicate significance difference from the corresponding control (Student’s t-test/ Mann-Whitney U-test, *P<0.05, n=3-4 animals/group).

Downstream intracellular signaling of IL-1 and IL-18 receptors (IL-1R1 and IL-18R1) includes recruitment of MYD88 (myeloid differentiation primary response 88),[14] for which mRNA expression was significantly upregulated in the both the amnion and chorion in the animals exposed to choriodecidual *Ureaplasma* infection (P<0.05, Fig. 4). In the chorion, NF-κB mRNA was also significantly higher in the *Ureaplasma* group compared to controls (P<0.05, Fig. 4B).

### Increased prostaglandin pathway proteins in fetal membranes of animals exposed to choriodecidual Ureaplasma infection

The constitutive form of cyclo-oxygenase, COX1/PTGS1 was identified in all fetal membrane samples and expression remained unchanged by infection (data not shown). An increase in *PTGS2* (inducible prostaglandin-endoperoxide synthase 2) gene expression (P<0.07) and a significant increase in *HPGD* (15-hydroxyprostaglandin dehydrogenase), involved in prostaglandin biosynthesis and degradation, respectively, were observed in the chorion samples collected from the animals in the *Ureaplasma* group compared to controls (Fig. 5B). In contrast, we observed significant decreases in both *PTGS2* and *HPGD* (P<0.05) gene expression in the amnion from animals in the infected group compared to control (Fig. 5A). However, by immunohistochemistry, PTGS2 protein staining was markedly higher in both the epithelial layer of the amnion and the chorion of fetal membranes from the *Ureaplasma* group when compared to the control group (Fig. 5C). Prostaglandin production was investigated in the amniotic fluid and no differences were identified in either PGE_2_ and PGF_2⍺_ concentrations between the control and *Ureaplasma* groups (Table 2).

**Figure 5.**
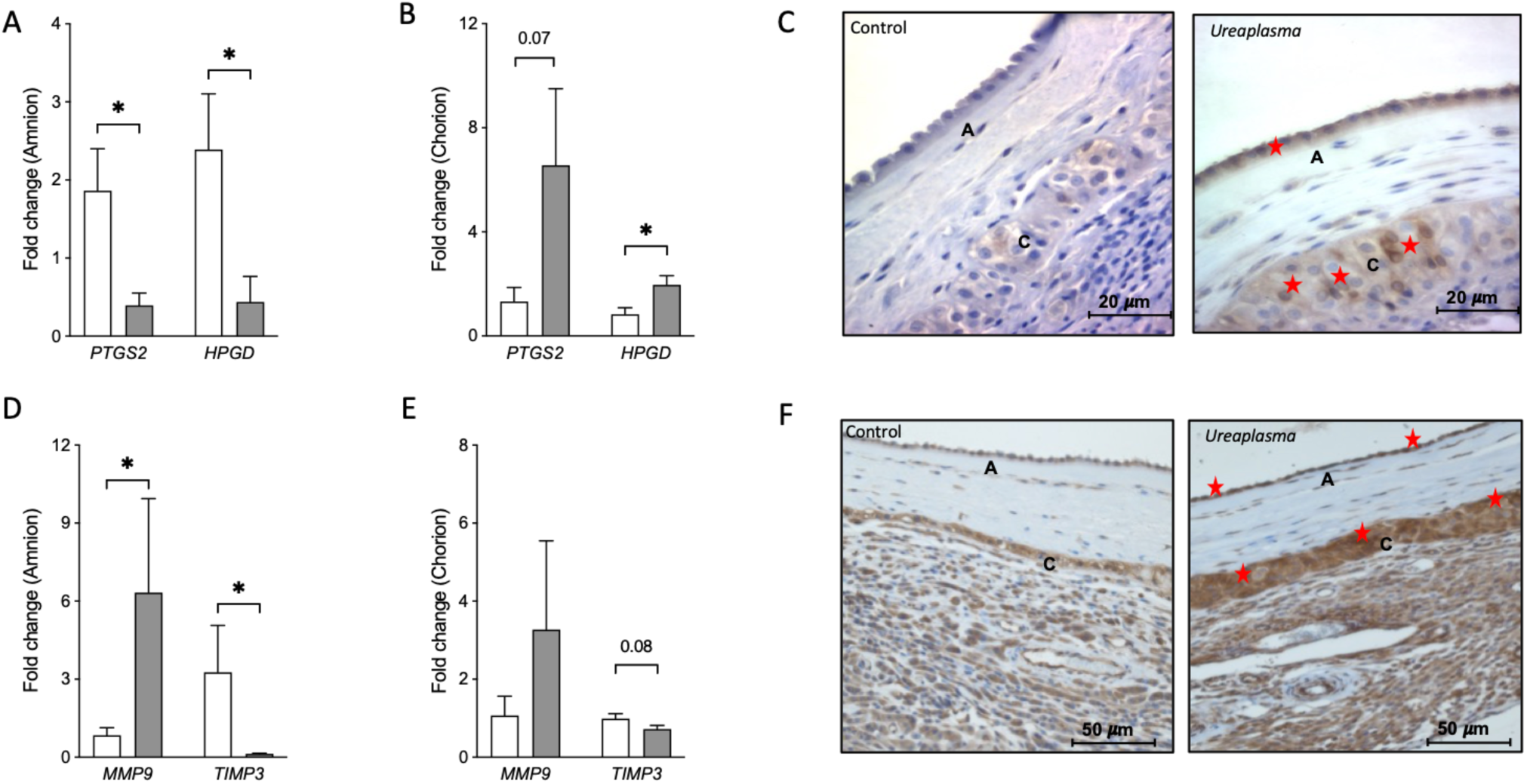
Choriodecidual *Ureaplasma parvum* infection induces gene and protein expression of the prostaglandin synthesizing enzyme PTGS2 and the collagenase MMP9 in chorioamnionic membranes. mRNA expression of *PTGS2* (Prostaglandin-endoperoxide synthase 2) and *HPGD (*15-hydroxyprostaglandin dehydrogenase) are shown in the (A) amnion and (B) chorion. mRNA expression for *MMP9* (matrix metallopeptidase 9) and *TIMP3* (tissue inhibitor of metalloproteases 3) are shown in the (D) amnion and (E) chorion. Representative immunohistochemical staining for (C) PTGS2 and (F) MMP9 for control and *Ureaplasma* animals is shown by brown staining (marked by stars) in membrane rolls including the amnion (a) and chorion (c). The results are expressed as fold changes in mRNA expression compared with control group and the housekeeping gene *GAPDH*. Individual bars represent mean±SEM fold change. Open bars are control and grey bars are the *Ureaplasma* group. The asterisks indicate significance difference from the corresponding control (Student’s t-test/ Mann-Whitney U-test, *P<0.05, n=3-4 animals/group).

### Increased level of enzymes capable of degrading extracellular matrix macromolecules in animals exposed to infection

An increase in gene expression of matrix metallopeptidase 9 (*MMP9*, P<0.05) with a corresponding decrease in MMP inhibiting protein, tissue inhibitors of metalloproteinases (*TIMP3*, P<0.05) was observed in the amnion (Fig. 5D). Neither MMP9 or TIMP3 gene expression in the chorion were significantly different between control and *Ureaplasma* animals (Fig. 5E). However, by immunohistochemistry, MMP9 protein localization was marked in both the epithelial layer of the amnion, and the chorionic fetal membranes collected from the *Ureaplasma* group compared to controls (Fig. 5F). Different MMP protein concentrations were also measured in the amniotic fluid and a significant decrease in the MMP-1 (P=0.04) and an increase in MMP-13 (P=0.04) was observed in animals from the *Ureaplasma* group (Table 2).

## DISCUSSION

Our results provide new experimental evidence that choriodecidual infection with *Ureaplasma parvum*, results in activation in the fetal membranes of inflammasome components and the induction of MMP collagenases and the prostaglandin synthesizing enzyme, PTGS2, without direct infection of the amniotic cavity. These findings suggest that even the early stages of intrauterine *Ureaplasma* infection initiates processes that can alter membrane structural integrity that are involved in the pathogenesis of pPROM and preterm labor. The unique, translational NHP model utilized in this study allows assessment of this early stage of ascending *Ureaplasma* infection, without microbial invasion of the amniotic cavity or preterm labor, providing a model of the initiation of preterm labor processes that are difficult to examine in other animal models (e.g., intra-amniotic or maternal inoculation) or clinical studies. These findings are particularly relevant to clinical cases of “sterile” intrauterine inflammation, when microbes are undetectable but inflammation is present.

A principle finding of our study was the upregulation of multiple genes that code for inflammasome (sensors, adaptor and effector) proteins and the associated IL-1 family inflammatory cytokines in response to choriodecidual *Ureaplasma* infection, despite the lack of MIAC or the onset of preterm labor in this model (see conceptual model, Fig. 6). The changes we identified included significantly increased NLRP3 mRNA and protein and NOD2 mRNA in the chorion, and significantly increased NLRC4 mRNA and AIM2 protein in the amnion. In addition, we identified increased mRNA expression for the adaptor protein, ASC, CASPASE-1 and IL-18, IL-1α and IL-1β. These findings suggest inflammasome activation by choriodecidual *Ureaplasma* infection. However, full activation, leading to processing of mature IL-18 and IL-1β protein expression in the membranes or amniotic fluid was not observed during this stage of ascending infection in our model of choriodecidual infection. Signaling via IL-1 and IL-18 receptors recruits intracellular MyD88 proteins (increased MyD88 in both chorion and amnion with *Ureaplasma* infection) which is upstream of various kinases which may lead to NFκB activation. NFκB, whose gene expression was also increased in the chorion with *Ureaplasma* infection, is a transcription factor classically associated with inflammation and involved in the pathophysiology of preterm labor.[15] NFκB plays multiple roles in inflammasome signaling, feedback and regulation of inflammasome mediated inflammation.[14] Increases in inflammasome sensor molecules and inflammasome activation have previously been reported in placental membranes in cases following ruptured membranes, from both pPROM and ruptured membranes at term.[16] Elevated concentrations of NLRP3, NLRC4 and NOD2 in the amniotic fluid are positively correlated with increased severity of amniotic fluid inflammation (sterile or infectious causes) and higher microbial burden in cases of chorioamnionitis, pPROM and spontaneous preterm labor.[17–20] In the current study, we identified activation of inflammasome components in the membranes without the need for microbial infection of the amniotic cavity, which may be representative of clinical cases of sterile intrauterine infection. Taken together with our findings, these previous studies support the hypothesis that the inflammasome activation that we identified during choriodecidual *Ureaplasma* infection may contribute to mechanisms of membrane weakening that could lead to pPROM.[21, 22] Activation of inflammasome mediated signaling pathways during the choriodecidual stage of ascending infection may therefore contribute to the pathogenesis of preterm labor caused by *Ureaplasma* infections.

**Figure 6.**
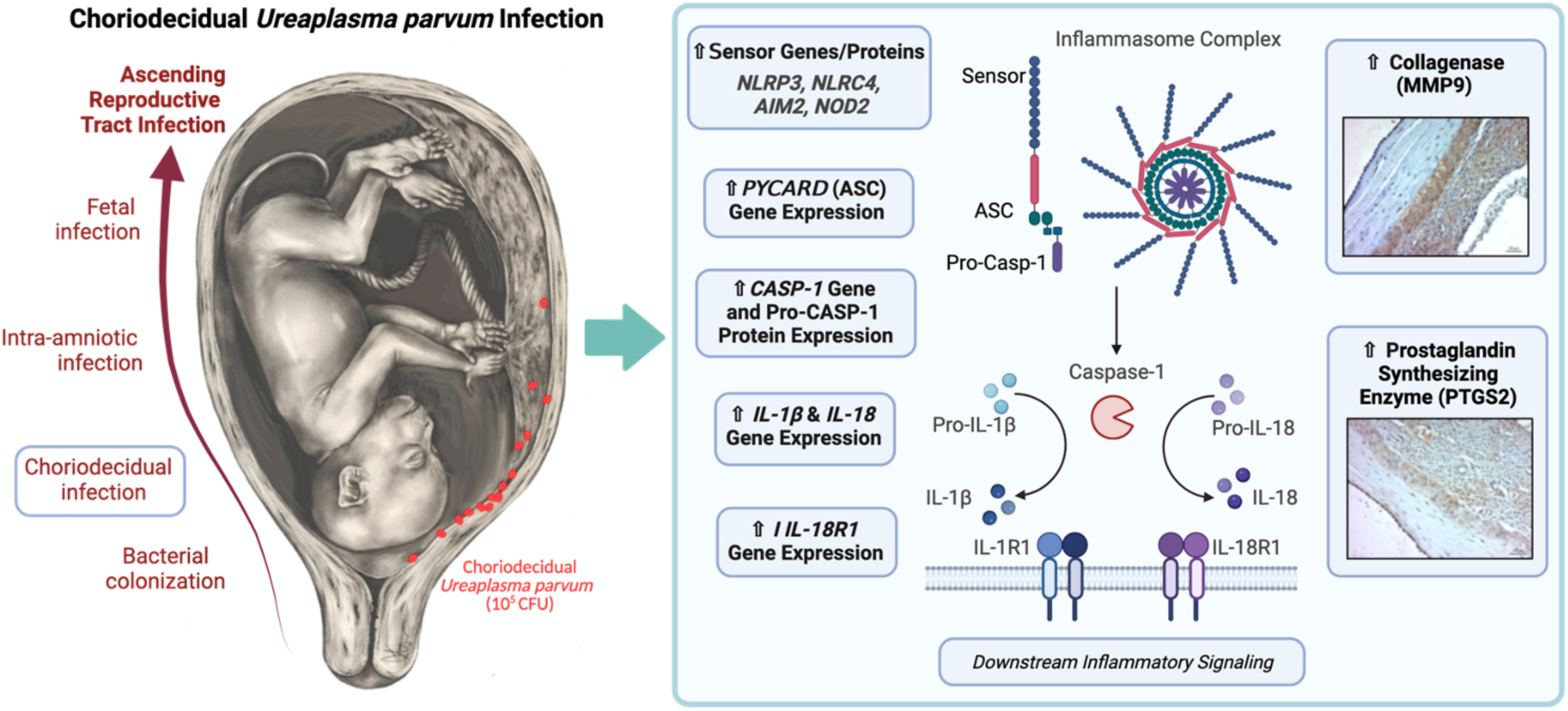
Conceptual model: Choriodecidual *Ureaplasma parvum* infection induces inflammasome activation in chorioamnionic membranes without amniotic fluid infection. Intrauterine infection and inflammation remain a significant cause of preterm labor and perinatal morbidity and mortality. Our findings suggest that *Ureaplasma parvum* infection can induce inflammasome activation *in vivo* in a clinically translational NHP model of intrauterine infection without microbial invasion of the amniotic cavity. Fetal membranes inflammasome activation likely activates CASPASE-1, which in turn converts the inactive pro-IL-1 *β* and IL-18 into mature IL-1 *β* and IL-18 forms. The mature IL-1 *β* and IL-18 propagates downstream inflammatory signaling that may aid in the further spread of microbes, progression of membrane weakening and initiation of pathological processes that ultimately lead to pPROM and preterm labor.

In addition to increases in multiple inflammasome sensing molecules in the current study, increased expression of ASC (*PYCARD*) mRNA in the chorion and *CASP-1* mRNA in both the amnion and chorion from pregnancies exposed to choriodecidual *Ureaplasma* infection was identified. We also observed an increase in immature pro-CASPASE-1 protein in the chorion. However, the lack of mature caspase-1 suggests that full inflammasome activation does not occur at this stage of choriodecidual *Ureaplasma* infection in this model. As the effector component of the inflammasome complexes, activation of CASPASE-1 is required for the cleavage of pro-IL-18 and pro-IL-1β into their active signaling forms.[7] The current study identified an upregulation in the expression of mRNA for IL-18, IL-1α and IL-1β genes in the amnion and IL-18 mRNA in the chorion. A concurrent increase in the IL-18 receptor (IL-18R1) and binding protein (IL-18BP), further demonstrates activation of this signaling pathway by *Ureaplasma* infection in the chorioamnionic membranes (Fig. 6). In previous studies in a non-human primate model, *Ureaplasma parvum* infused directly into the amniotic fluid, produces a robust inflammatory response, including increased IL-1β concentrations in the amniotic fluid (although IL-18 concentrations were not reported), that resulted in preterm labor and fetal pneumonia.[3] In a similar NHP study, direct intra-amniotic infusion of IL-1β, also resulted in preterm labor.[23] Studies of urinary tract infections and infertility studies in males have previously reported elevated IL-1β and IL-18 concentrations in urine and semen, respectively, in response to urogenital infection with *Ureaplasma parvum.*[24, 25] *Nlrp3* knockout studies in mice have demonstrated that IL-1β production in the amniotic cavity in response to LPS occurs in both Nlrp3 dependent and independent pathways. [26] Both IL-1α and IL-1β have also previously been associated with preterm labor, chorioamnionitis and intrauterine inflammasome activation. [27–29] As a member of the IL-1 family, increased IL-18 in the amniotic fluid is also correlated with microbial infection of the amniotic cavity and the onset of preterm labor. [30–32] IL-18 has also previously been linked with pPROM.[33] However, despite increased gene expression, increases in the cleaved active forms of IL-1β or IL-18 were not observed. This suggests that inflammasome activation is initiated by choriodecidual *Ureaplasma* infection in this model but that production of active IL-1β and IL-18 is limited at this early stage of infection. This may explain why in our model, we did not identify increases in amniotic fluid IL-1 β and IL-18 that is likely required for the onset of preterm labor. We hypothesize that progression of infection to MIAC and subsequent increase in bacterial load may be required for full inflammasome activation that results in inflammation sufficient to drive preterm labor. Despite choriodecidual *Ureaplasma* infection does not result in preterm labor or significant increases in amniotic fluid pro-inflammatory mediators in this model, the upregulation of inflammatory genes in the membranes demonstrate that this early stage of ascending infection may be important for the initial priming of processes that result in pPROM and eventually preterm labor.

MMP9 is the major collagenase enzyme involved in both physiological and pathological labor and has a significant role in membrane degradation and rupture.[34] In our study, the expression of MMP9 mRNA and protein were markedly increased in both the amnion and chorion from animals exposed to choriodecidual *Ureaplasma* infection. MMP9 expression has previously been associated with intrauterine Ureaplasma infection and in membrane explants.[3, 35] Increased MMP-9 amniotic fluid concentrations in cases of spontaneous preterm birth that were accompanied by positive amniotic fluid *Ureaplasma* cultures, compared to preterm birth cases negative for *Ureaplasma*.[36] Human neutrophils cultured with *Ureaplasma* also demonstrated an increase in MMP9 activity,[36] showing MMP9 expression as part of the immune response to *Ureaplasma* infection. *In vitro* studies have also shown MMP9 production in response to *Ureaplasma* infection in both adult and neonatal monocytes.[37] Along with the increase of *MMP9* mRNA in the amnion, we also identified a concurrent decrease in *TIMP3* mRNA. TIMP3 is a metallopeptidase inhibitor and its downregulation likely contributes to MMP9 action on membrane collagen degradation. *In vivo* cultures of human fetal membranes have identified similar increases in MMP9, coupled with a synergistic reduction in TIMP3, and in association with markers of membrane weakening when cultured with the cytokines IL-1 β and TNF*α*.[38] Both IL-1 β and TNF*α* gene expression was also increased in the membranes of the current study. This study therefore demonstrates activation of MMP9 collagenase pathways that are well-known to perturb membrane integrity leading to membrane rupture and pPROM. The upregulation of these pathways before *Ureaplasma* infection is detectable in the amniotic fluid, suggests that membrane integrity may be affected prior to MIAC and is a factor that may mediate passage of microbes across the membranes to invade the amniotic cavity. This is a significant finding since the mechanisms of *Ureaplasma* transmigration to invade the amniotic cavity are currently poorly understood.

PTGS2 is also well-studied in terms of its role in responding to intrauterine infection by producing pro-inflammatory prostaglandins that contribute to uterine contractility and cervical ripening. Two pro-inflammatory cytokines that demonstrated increased gene expression in the amnion and chorion, respectively, IL-1*β* and TNF*α*, have both been shown to increase NOD2 expression *in vitro* when incubated with fetal membrane explants. [39] In the same study, stimulation of membrane explants with a NOD2 ligand also stimulated increased PTGS2 gene expression and increased MMP9 enzyme activity. Similar increases were also identified in our *in vivo* model of choriodecidual *Ureaplasma* infection. Taken together with inflammasome activation and upregulation of inflammatory genes in the fetal membranes in this model, increased MMP9 and PTGS2 suggest that mechanisms of membrane weakening and preterm labor are initiated even at this early stage of ascending infection by *Ureaplasma*, thus providing insight into the onset of these processes during a timepoint for which we currently have little data.

We have shown in this study, that choriodecidual *U. parvum* inoculation results in infection that can remain localized within the choriodecidual membranes, even with up to 20 days of infection. We hypothesize that the relatively low dose of inoculum may be too low bacterial load to passage across membranes. Previous *in vitro* studies comparing *Ureaplasma* exposure on either the choriodecidual or amnionic side of membrane explants in a dual-chamber culture system, showed that increases in bacterial dose on the choriodecidual side was not reflected with a concurrent increase of passage of *Ureaplasma* to the amniotic compartment. Suggesting an unknown mechanism may be involved in the passage of *Ureaplasma* across intact membranes, resulting in MIAC. The lack of labor-like contractions and minimal production of pro-inflammatory mediators in the amniotic fluid with choriodecidual infection in the current study supports that the inflammatory stimuli or bacterial load with localized choriodecidual infection was not sufficient to cause MIAC and subsequently produce preterm labor. The mechanisms needed for infection progress from the membranes to the amniotic cavity still need to be elucidated. Sufficient microbial burden, timing and specific location of the infection, gestational age, individual genetic and epigenetic vulnerability and polymicrobial infection have been postulated as factors that would influence the ability of *Ureaplasma* microbes to invade the amniotic cavity. *Ureaplasma* spp. are known for their relatively low virulence and ability to produce a prolonged, low-grade infection, potentially due to immunomodulatory properties.[4, 40] Unlike bacterial infections, or experimental models such as those that utilize LPS, that result in exponential production of intrauterine pro-inflammatory mediators and precipitous preterm labor, *Ureaplasma* infection has been detected for extended periods in the amniotic fluid before the onset of labor, which may contribute to their role in preterm labor and neonatal morbidity and explain why not all women colonized with *Ureaplasma parvum* progress to intra-uterine infection or preterm labor. We hypothesize that the membrane inflammasome activation identified in the current study is a mechanism that may be integral in the loss of membrane cytostructural integrity that could contribute to the passage of microbes across intact membranes into the amniotic cavity, along with other mechanisms not included in the current study. Additionally, activation of the pathways identified in this study, would likely be involved in processes leading to the premature rupture of membranes. There is a known link between inflammasome activation and pPROM, between MMP9 and PTGS2 expression and pPROM and strong clinical association between pPROM positive amniotic fluid cultures for *Ureaplasma*.[36, 41]

Advantages of the current study include the use of a translational NHP model and live bacterial infection with a low passaged clinical isolate of *Ureaplasma parvum* as the infectious stimuli. In addition, the use of our established chronically catheterized pregnant rhesus monkey model allows us to investigate the mechanisms of *Ureaplasma* infection isolated to the choriodecidual junction and compare these findings to previous studies of intra-amniotic inoculation. This location of infection that would be difficult to study in other *in vitro* or *in vivo* models. Catheterization also allows real-time monitoring of uterine contractility and serial sampling of amniotic fluid and maternal blood over the course of the experiment. A disadvantage of the current study, was that we did not include measurement of inflammasome complex proteins in the amniotic fluid, which may have provided further information about whether inflammasome activation had spread further than the fetal membranes. Future studies will aim to identify whether downstream inflammatory pathways, including NFκB, are activated during this stage of infection, or whether location or a threshold of bacterial burden must be reached to trigger preterm labor. However, inflammation present in the fetal membranes indicates a potential source of inflammation that may be able to reach the fetus. Activation of the fetal inflammatory response during the early stages of ascending reproductive tract infection may contribute to vulnerability to perinatal injury *in utero* or following premature delivery.

Inflammasome-mediated inflammation resulting from *Ureaplasma* infection may also be a potential target for treatment to prevent preterm labor. Azithromycin(AZI) has specific effects on reducing NLRP3 mediated inflammasome inflammation, including in neonatal blood monocytes.[42–45] Previous NHP studies with intra-amniotic *Ureaplasma* infection has identified azithromycin as an effective treatment in delating preterm labor and improving fetal lung inflammation. Further investigation of the role of AZI in inhibiting inflammasome-mediated preterm labor processes may improve our understanding and development of therapies to prevent preterm labor, including recent interest in the inhibition of IL-1 *β* pathways which are present in the pathways identified in the current study.

This study therefore demonstrates that *Ureaplasma parvum* infection can induce inflammasome activation *in vivo,* predominantly at the mRNA level, in a clinically translational NHP model of intrauterine infection and that this activation can occur despite lack of widespread infection of the amniotic cavity. Activation of inflammasome complexes in the chorioamnionic membranes during an early stage of ascending reproductive tract infection may be an important step in the proliferation of intrauterine infection and inflammation. This model is particularly relevant to clinical cases of sterile intrauterine inflammation in which microbes are undetectable in the amniotic fluid. Development of this NHP model of choriodecidual *Ureaplasma* infection will therefore aid future studies aimed at elucidating how transmigration of *Ureaplasma* microbes across the chorioamnionic membranes to invade the amniotic cavity can occur, the mechanisms of which remains poorly understood. These translational NHP studies are aimed at investigating the processes that initiate pPROM and preterm labor that may be novel targets for diagnostic tests and therapeutic approaches to prevent preterm birth and improve neonatal and long-term outcomes for premature infants that are exposed to inflammation during gestation.

## Acknowledgments

We would like to acknowledge the Oregon National Primate Research Center, Animal Resources & Research Support Unit and specifically the Surgical Services Unit, Dr. Drew Martin, DVM and Darla Jacobs for their project support. Research reported in this publication was supported by the Eunice Kennedy Shriver National Institute of Child Health and Human Development, R00HD090229 and the Office of the Director, National Institutes of Health under Award Number P51OD011092 to the Oregon National Primate Research Center. The content is solely the responsibility of the authors and does not necessarily represent the official views of the National Institutes of Health.

