## Supplemental Figures for "Membrane Inflammasome Activation by Choriodecidual *Ureaplasma parvum* Infection without Intra-Amniotic Infection in an NHP Model"

**Supplementary Table 1:** *Ureaplasma* Culture and PCR Results for Vehicle Control and *Ureaplasma* inoculated Groups

| | **Control** | | ***Ureaplasma*** | | | | | | | |
|---|---|---|---|---|---|---|---|---|---|---|
| | All Controls (1-4) | | U.p. 5 | | U.p. 6 | | U.p. 7 | | U.p. 8 | |
| | CFU/mL | PCR | CFU/mL | PCR | CFU/mL | PCR | CFU/mL | PCR | CFU/mL | PCR |
| Pre-Inoculation Amniotic Fluid | 0 | - | 0 | - | 0 | - | 0 | - | 0 | - |
| Post-Inoculation Amniotic Fluid | 0 | - | 0 | - | 0 | - | 0 | - | 0 | - |
| Membranes at Inoculation Site | 0 | - | 150 | + | 1.5 x 10^5 | + | 1500 | + | 1.3 x10^3 | + |
| Membranes (distal) | 0 | - | 0 | + | 500 | + | 0 | - | 100 | + |
| Placenta Tissue Sample | 0 | - | 0 | - | 1600 | + | 0 | - | 1 | + |

**Supplementary Table 2:** List of RT-qPCR Primers for Macaca Mulatta

| Primer # | Gene symbol | Gene name | Primer sequences | |
|---|---|---|---|---|
| | | | FP/RP | Sequences (5' to 3') |
| 1 | *NLRP3* | NACHT, LRR and PYD domains containing protein 3 | FP | CAGACTCTACGTGGGGGAGA |
| | | | RP | GCCAGAATTCACCAACCCCA |
| 2 | *NLRC4* | NLR family CARD domains containing 4 | FP | TTCAGGACTTGAATGGACAAAGTCT |
| | | | RP | GTGTTGGTCCTTCCTCCACA |
| 3 | *AIM2* | Absent in melanoma 2 | FP | CGGAGGCAACCTGAAGGTGA |
| | | | RP | GACGACTTTGGGATCAACATTCC |
| 4 | *NOD2* | Nucleotide binding oligomerization domain containing 2 | FP | GGTGTCTGCAAGGCTCTGTA |
| | | | RP | GTGGTTATTCCCCAGCCTCAA |
| 5 | *PYCARD* | PYD and CARD domain containing | FP | AGCTGGTCAGCTTCTACCTG |
| | | | RP | TCCAGAGCCCTGGTGC |
| 6 | *CASP1* | Caspase-1/ Interleukin-1 converting enzyme | FP | ATGCCCACCACTGAAAGAGTG |
| | | | RP | GGATCTCTTCACTTCCTGCCCA |
| 7 | *CASP4* | Caspase-4 | FP | CTGTTAACTATGGCGGAGGGC |
| | | | RP | GAGTCTGCCATGACCCGAAC |
| 8 | *TNF-α* | Tumor necrosis factor-alpha | FP | TCTTCTCCTTCCTGCTCGTGG |
| | | | RP | TCAGCTTGAGGGTTTGCTACAAC |
| 9 | *IL-6* | Interleukin-6 | FP | TCCTGCAGAAAAAGGCAAAGAA |
| | | | RP | AAGCTGCGCAGGATGAGATG |
| 10 | *IL-8* | Interleukin-8 | FP | CGGAAGGAACCATCTCGCTC |
| | | | RP | GGCAAAACTGCACCTTCACAC |
| 11 | *IL-1α* | Interleukin-1 alpha | FP | CCAAGCTGACCTTCAAGCAG |
| | | | RP | CTGGGCTTGATGATTTCTTCCTC |
| 12 | *IL-1β* | Interleukin-1 beta | FP | GATGGCTTACTACAGCGGCA |
| | | | RP | AAGCCCTCGTTGTAGTGCTC |
| 13 | *IL-1RN* | Interleukin-1 receptor antagonist | FP | AGTCTGGAAGACCTCGGAAG |
| | | | RP | CTGGTTAACATCCCAGATTCTGAAG |
| 14 | *IL1R1* | Interleukin 1 receptor type 1 | FP | GGATGAACGGAGATGGTCCAG |
| | | | RP | ACTCAATTGTCCAGGCCAGC |
| 15 | *IL-18* | Interleukin-18 | FP | GACAGTACGCTTTACTTTATAGCTG |
| | | | RP | GTCCGGGGTGCATTATCTCT |
| 16 | *IL-18BP* | Interleukin 18 binding protein | FP | AGTGCGCTCGGTGTATTTCC |
| | | | RP | CCTTCACCTTCACACTGGCTTC |
| 17 | *IL-18R1* | Interleukin-18 receptor 1 | FP | GGATCGCTGCTTCTCACCTA |
| | | | RP | GCTTCAAACGGCTTCTCTTCA |
| 18 | *PTGS2* | Prostaglandin-endoperoxide synthase 2 | FP | TCCCTTGGGTGTGAAAGGTAAG |
| | | | RP | CAGCCCTTTGGTGAAAGCTG |
| 19 | *HPGD* | 15-hydroxyprostaglandin dehydrogenase | FP | ATGTCATCTTTGGCAGGACTCA |
| | | | RP | ACAAAGCCTGGACAAATGGC |
| 20 | *MMP9* | Matrix metallopeptidase 9 | FP | TTCTGCCCGGACCAAGGATA |
| | | | RP | ACATAGGTACATGAGCGCC |
| 21 | *TIMP3* | TIMP metallopeptidase inhibitor 3 | FP | TCAAGCAGATGAAGATGTACCG |
| | | | RP | TGGTACTTGTTGACCTCCAGC |
| 22 | *GAPDH* | Glyceraldehyde 3-phosphate dehydrogenase | FP | GAAATCCCATCACCATCTTCCAGG |
| | | | RP | GAGCCCCAGCCTTCTCCATG |
| 23 | *MYD88* | Myeloid differentiation primary response 88 | FP | CATCGAAAAGAGGTGCCGC |
| | | | RP | CAGGGGTTGGTGTAATCGCA |
| 24 | *NF-κB1* | Nuclear factor kappa B subunit 1 | FP | CTCGCCACCAGGCTTCAG |
| | | | RP | GCAGTGCCATCTGTGGTTG |

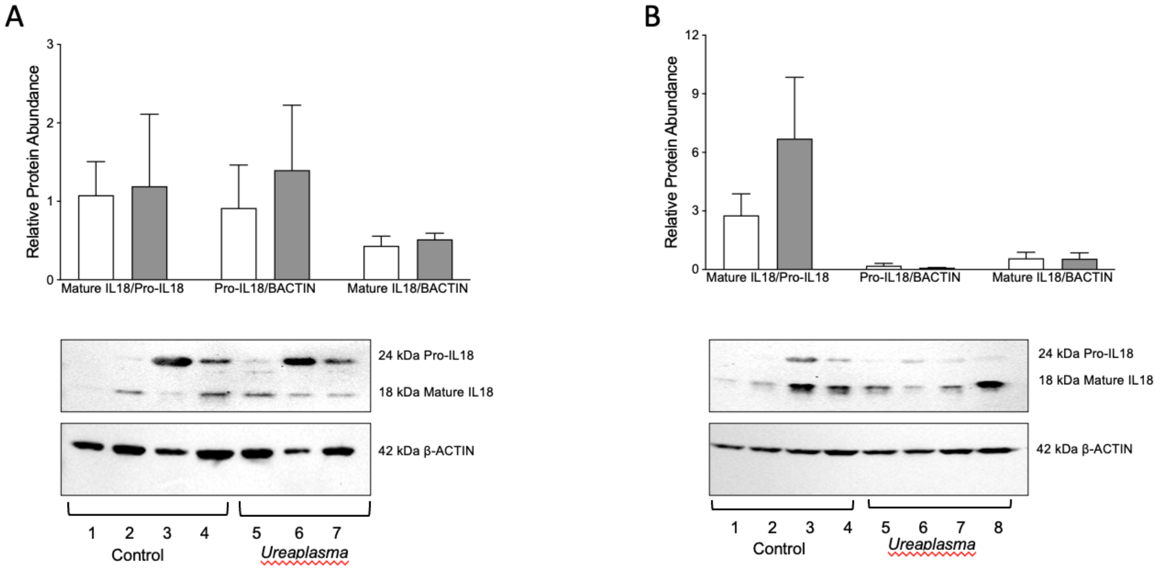


**Supplementary Figure 2. Choriodecidual *Ureaplasma parvum* infection did not show a significant change in the levels of mature IL-18 protein in the chorioamnionic membranes.** Densitometric quantification and representative western blot of pro- and mature form of IL-18 in the (A) amnion and (B) chorion from control (open bars) and *Ureaplasma (U.p,,* grey bars*)* group, as indicated. $\beta$-actin serves as a loading control. The results are shown as relative protein levels with respective to the loading control, and compared with control group. Each bar represents the mean±SEM (n=3-4 animals/group).

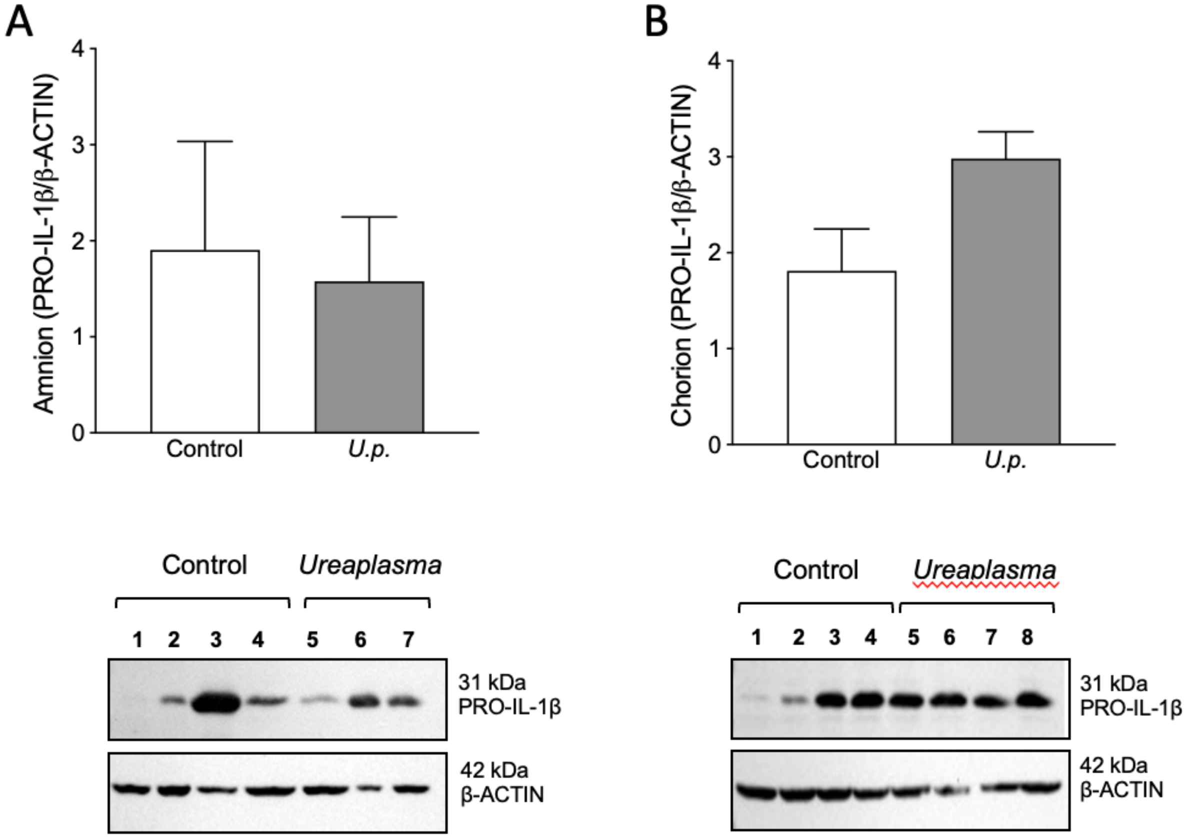


**Supplementary Figure 3. Choriodecidual *Ureaplasma parvum* infection did not show a significant change in the levels of IL-1β protein in the chorioamnionic membranes.** Densitometric quantification and representative western blot of pro-IL-1$\beta$ in the (A) amnion and (B) chorion from control (open bars) and *Ureaplasma (U.p,,* grey bars*)* group, as indicated. $\beta$-actin serves as a loading control. The results are shown as relative protein levels with respective to the loading control, and compared with control group. Each bar represents the mean±SEM (n=3-4 animals/group).

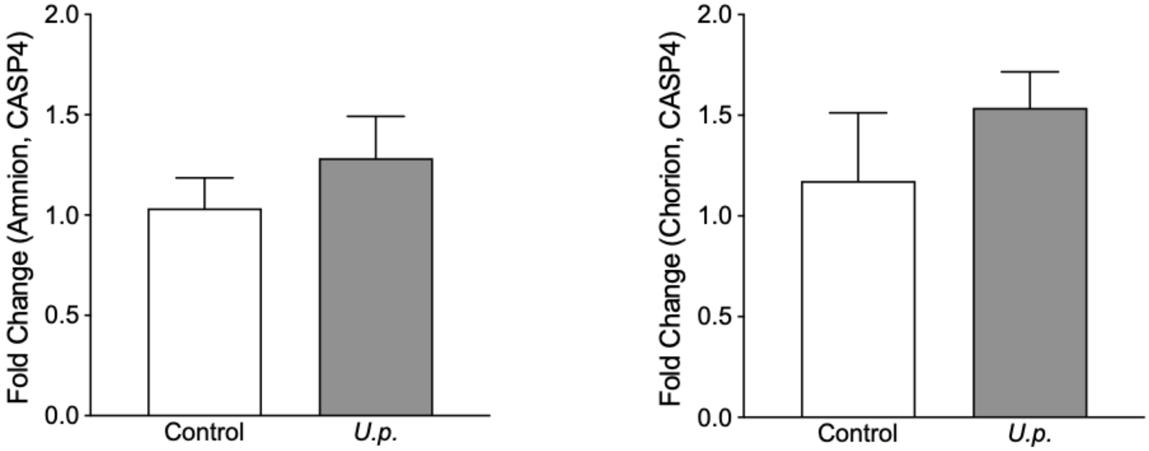


**Supplementary Figure 3:** q(RT)PCR expression of *CASP-4* in the (A) amnion and (B) chorion collected during cesarean section delivery. The results are expressed as fold changes in mRNA expression compared with control group and the housekeeping gene *GAPDH*. Individual bars represent mean±SEM fold change. Open bars are control and grey bars are the *Ureaplasma* group. (n=3-4 animals/group).
